# Statin treatment and the risk of depression

**DOI:** 10.1101/349332

**Authors:** Ole Köhler-Forsberg, Christiane Gasse, Liselotte Petersen, Andrew A. Nierenberg, Ole Mors, Søren Dinesen Østergaard

## Abstract

The effect of statin treatment on the risk of developing depression remains unclear. Therefore, we aimed to assess the association between statin treatment and depression in a nationwide register-based cohort study with up to 20 years of follow up. We identified all statin users among all individuals born in Denmark between 1920 and 1983. One non-user was matched to each statin user based on age, sex and a propensity score taking several potential confounders into account. Using Cox regression we investigated the association between statin use and: I) redemption of prescriptions for antidepressants, II) redemption of prescriptions for any other drug, III) depression diagnosed at psychiatric hospitals, IV) cardiovascular mortality and V) all-cause mortality. A total of 193,977 statin users and 193,977 non-users were followed for 2,621,282 person-years. Statin use was associated with I) increased risk of antidepressant use (hazard rate ratio (HRR)=1.33; 95% confidence interval (95%-CI)=1.31-1.36), II) increased risk of any other prescription drug use (HRR=1.33; 95%-CI=1.31-1.35), III) increased risk of receiving a depression diagnosis (HRR=1.22, 95%-CI=1.12-1.32) - but not after adjusting for antidepressant use (HRR=1.07, 95%-CI=0.99- 1.15), IV) reduced cardiovascular mortality (HRR=0.92, 95%-CI=0.87-0.97) and V) reduced all-cause mortality (HRR=0.90, 95%-CI=0.88-0.92). These results suggest that the association between statin treatment and antidepressant use was unspecific (equivalent association between statins and other drugs) and that the association between statin use and depression diagnoses was mediated by antidepressant use. Thus, statin users and non-users appear to be equally likely to develop depression, but the depression is more often detected/treated among statin users.

## INTRODUCTION

Statins (HMG-CoA reductase inhibitors) are widely used in the primary and secondary prevention of cardiovascular disease (1–5). In addition to lowering cholesterol, statins have anti-inflammatory effects independent of their cholesterol-lowering activity (6). Since inflammation is considered to play an important role in the pathophysiology of depression (7–9) it has been investigated whether the combination of treatment with a statin and a selective serotonin reuptake-inhibitor (SSRI) has superior antidepressant effect compared to an SSRI alone - a standard first line pharmacological treatment for depression (10,11). Indeed, the results from three small, randomized controlled trials (RCTs) have shown that the combination of a statin and an SSRI has superior antidepressant effect compared to treatment with an SSRI alone (12). We recently investigated whether this effect could be generalized to the population-based level via a nationwide cohort study of 872,216 incident SSRI users of whom 113,108 used a statin concomitantly. We found that the combination of treatment with a statin and an SSRI was associated with an approximately 30% (and statistically significant) reduction in the risk of admission for depression compared to treatment with an SSRI alone (13).

Due to the possible synergistic antidepressant effect of a statin combined with an SSRI, the potential protective effect of statins in relation to depression onset is of considerable interest. Many studies of statins, both RCTs (5,14,15) and observational studies (16–20) have investigated this aspect either as primary and secondary outcome, and both increased and decreased risks with statin treatment have been detected. Thus, the effect of statin treatment on the risk of depression remains unclear.

In the study presented here, we investigated the effect of statin treatment on the risk of incident depression by means of a propensity-score matched analysis based on longitudinal data from nationwide Danish registers. Propensity-score matching is a statistical method used to mitigate confounding by indication in observational studies (21). The propensity score represents the probability of an individual receiving a treatment given this individual’s pre-treatment characteristics (age, sex, morbidity, etc.). The purpose of propensity score matching is to generate a control group that is as close to identical to the treatment group on the pre-treatment characteristics as possible, thereby mimicking the similarity of groups in RCTs, who are assumed to only differ in terms of the treatment they have been randomly allocated to (21).

## METHODS

### Design

This is a historical prospective propensity-score matched cohort study based on data from nationwide Danish registers. Linkage of register data at the level of the individual was possible due to the unique personal registration number, which is assigned to all Danes at the time of birth and registered in the Danish Civil Registration System (22). The design of the study in terms of definition of population and matching procedures is described below and illustrated in Figure 1.

**Figure 1.**
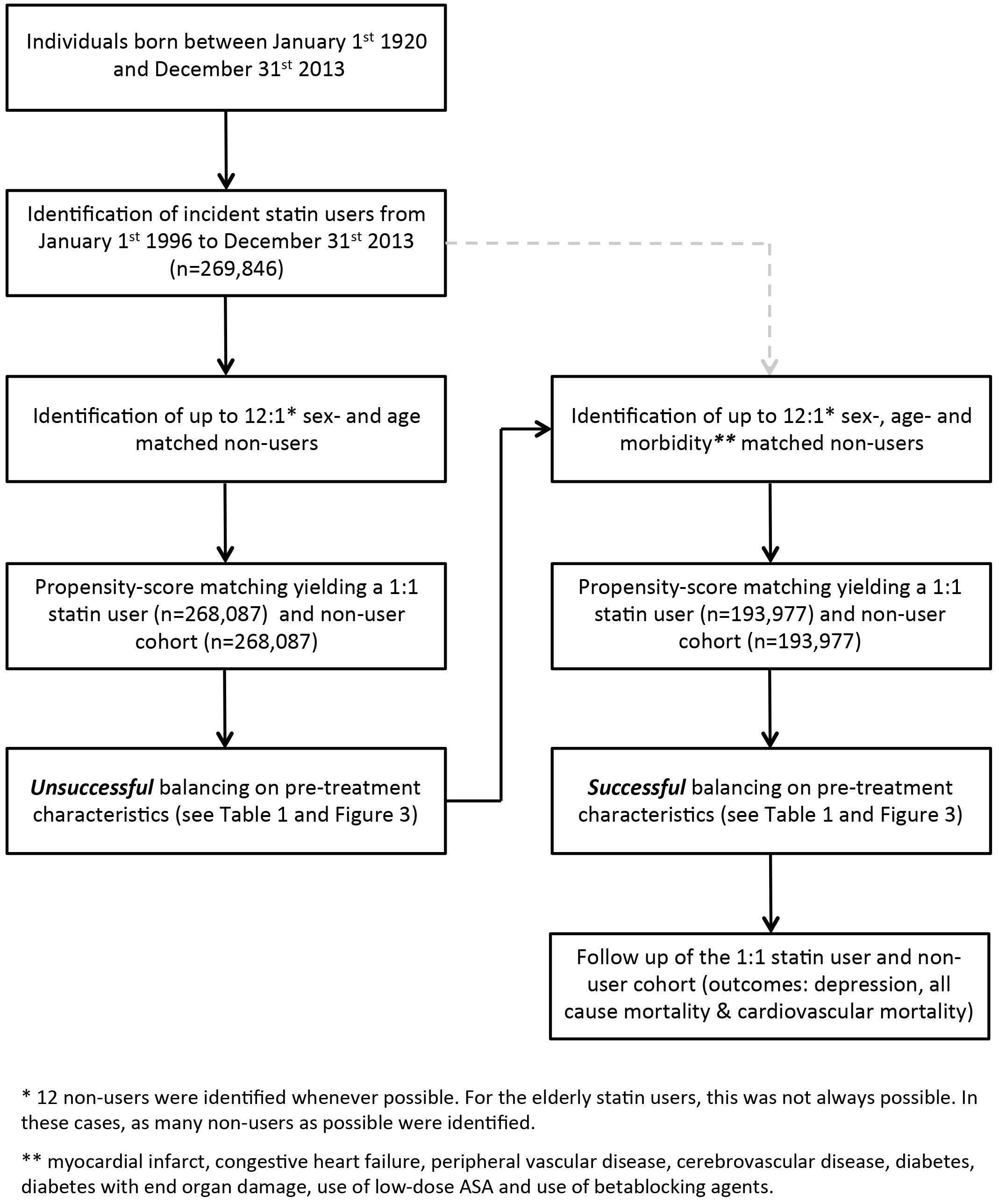
Overview of the matching of non-users to statin users

### Population-based sample

Individuals treated with statins and propensity-score matched controls were identified among all individuals born in Denmark between January, 1, 1920, and December, 31, 1983, who were alive on their 30^th^ birthday and on January, 1, 1996 identified via the Danish Civil Registration System (22).

### Definition of incident statin users

Individuals aged ≥30 years redeeming a prescription for a statin (Anatomic Therapeutic Chemical (ATC) code: C10AA) between January, 1, 1996 and December, 31, 2013 were defined as statin users. The index date was the date of redemption of a statin prescription. Information on redeemed prescriptions was extracted from the Danish National Prescription Registry (DNPR) (23), which contains data on all prescriptions redeemed at Danish pharmacies since January 1, 1995. To obtain a cohort of incident statin users, we required that the statin users had not redeemed a prescription for a statin in the year prior to the index date. We also excluded individuals who were registered with a redeemed prescription for a statin prior to the age of 30. In order to study incident depression and to rule out prior severe psychopathology in the study population, we required that the statin users had not redeemed a prescription for one of the following psychopharmaceuticals within the year prior to their index date: antidepressants (ATC-code: N06A), antipsychotics (N05A excl. N05AN), lithium (N05AN), or other mood-stabilizers (oxcarbazepine, carbamazepine, lamotrigine, valproate, topiramate, gabapentin, phenytoin (N03AF02, N03AF01, N03AX09, N03AG01, N03AX11, N03AX12, N03AB02)). Finally, individuals registered with a diagnosis of depression (International Classification of Diseases, 8^th^ revision (24) (ICD-8) 296.09, 296.29, 298.09, 300.49 / International Classification of Diseases, 10^th^ revision (25) (ICD-10) F32, F33), bipolar disorder (ICD-8: 296.19, 296.39, 298.19 / ICD-10: F30, F31, F38.00), or schizophrenia / schizoaffective disorder (ICD-8: 295.×9, 296.89 / ICD-10: F20, F25) in the Danish Psychiatric Central Research Register (DPCRR) between January, 1, 1969 and their index date were not eligible for the study. The DPCRR contains information regarding all diagnoses assigned in relation to admissions to psychiatric hospitals in Denmark since 1969 as well as diagnoses assigned following outpatient treatment and emergency room contacts since 1995 (26,27)Via this procedure we identified 269,846 incident statin users. Subsequently, we applied a stepwise matching procedure including propensity score matching, equivalent to that of Nielsen et al. (28).

### Sex- and age matching of non-users to each statin user (matching step 1)

For each statin user, we randomly drew up to 12 sex and age (month of birth) matched controls, who had not redeemed a statin prescription on the index date of the statin user, nor in the previous year. Hence, among the matched non-users, the index date was the same date as the index date of the statin users. For statin users for whom it was not possible to match 12 non-users on sex and age (predominantly among the eldest statin users), as many matched non-users as possible were included. The eligibility criteria in terms of psychiatric morbidity and use of psychopharmaceuticals described for the statin users were also employed in the selection of the non-users. An individual initially included as a non-user could be included in another matched pair as a statin user if redeeming a prescription for a statin. Similarly, an individual initially included as a statin user could be included in another matched pair as a non-user if this individual had not redeemed a statin prescription within the year prior to the new matched date.

### Propensity score matching (matching step 2)

As a second matching step, we calculated the propensity score for each individual using the pre-treatment characteristics we considered in our recent study on the effect of concomitant treatment with an SSRI and a statin as covariates (including indications for statin use and a range of other characteristics associated with both statin use and depression) (13). Specifically, for each individual, we extracted data on the following covariates at their index date: educational level (primary school, secondary school or higher education) (29); the 19 medical conditions from the Charlson Comorbidity Index (30) modified for use with the ICD-10 (31): myocardial infarct, congestive heart failure, peripheral vascular disease, cerebrovascular disease, dementia, chronic pulmonary disease, connective tissue disease, ulcer disease, mild liver disease, diabetes, hemiplegia, moderate to severe renal disease, diabetes with end organ damage, any tumor, leukemia, lymphoma, moderate to severe liver disease, metastatic solid tumor, and AIDS - all dichotomously defined based on main diagnoses registered in the Danish National Patient Register, which covers all diagnoses assigned at hospitals since 1977 (32); use of the following medications (dichotomously defined based on redeemed prescriptions within the year preceding the index date): corticosteroids (ATC-codes: H02A and H02B), anti-inflammatory/anti-rheumatic agents (M01B and M01C), non-steroidal antiinflammatory drugs (NSAIDs; M01A and N02BA), low-dose acetylsalicylic acid (B01AC06), antihypertensives (C02), diuretics (C03), peripheral vasodilators (C04), vasoprotective agents (C05), betablocking agents (C07), calcium channel blockers (C08), agents acting on the renin-angiotension system (C09), lipid modifying agents other than statins (C10AB, C10AC, C10AD, C10AX), anxiolytics (N05BA, N05CD02, N05CD05, N05CD06, N03AX16), and benzodiazepine-like hypnotics (N05CF01, N05CF02).

Covariates to be included in the propensity score are those associated both with exposure and with outcome (or at least with outcome) (21). Therefore, we evaluated whether the individual covariates were associated with both the exposure (statin treatment) and the individual outcomes considered in the study, i.e., I) redemption of a prescription for an antidepressant, II) redemption of a prescription for any other drug, III) depression diagnosed at psychiatric hospitals, IV) cardiovascular disease, V) cardiovascular mortality and VI) all-cause mortality) using logistic regression. These analyses showed that all covariates were associated with both the exposure (statin use) and all the outcomes, except for the following covariates, which were not associated with some specific outcomes: a) AIDS was not associated with redemption of a prescription for an antidepressant; b) moderate to severe liver disease, AIDS, and use of peripheral vasodilators were not associated with redemption of a prescription for any other drug; c) peripheral vascular disease, diabetes, diabetes with end organ damage, leukemia, and lymphoma were not associated with depression diagnosed at psychiatric hospitals; d) AIDS was not associated with cardiovascular disease; e) leukemia, lymphoma, and use of vasoprotective agents was not associated with cardiovascular mortality; and f) AIDS was not associated with all-cause mortality. Consequently, we performed the propensity score matching procedure both with and without including these covariates for each specific outcome, to see if this difference had any impact on the risk estimates. This was not the case and we therefore report results from the analyses in which all covariates have been used to calculate the propensity score.

In order to obtain the propensity score, we calculated a logistic regression model for each individual based on the covariates outlined above. For each statin user, we then chose the non-user (1:1 matching among the up to 12 matched individuals) with the propensity score closest to that of the statin user by means of the nearest neighbour matching algorithm. Via this procedure, we were able to match 268,087 non-users to 268,087 of the 269,846 statin users. Subsequently, we tested the balance of the pre-treatment covariates between the 268,087 statin users and the 268,087 propensity score matched non-users by means of standardized differences. An absolute standardised difference <10% and a variance ratio between 0.8 and 1.25 is generally considered to support the assumption of balance between groups (33,34). The balance between the statin users and non-users was also assessed graphically (See Figure 2).

**Figure 2.**
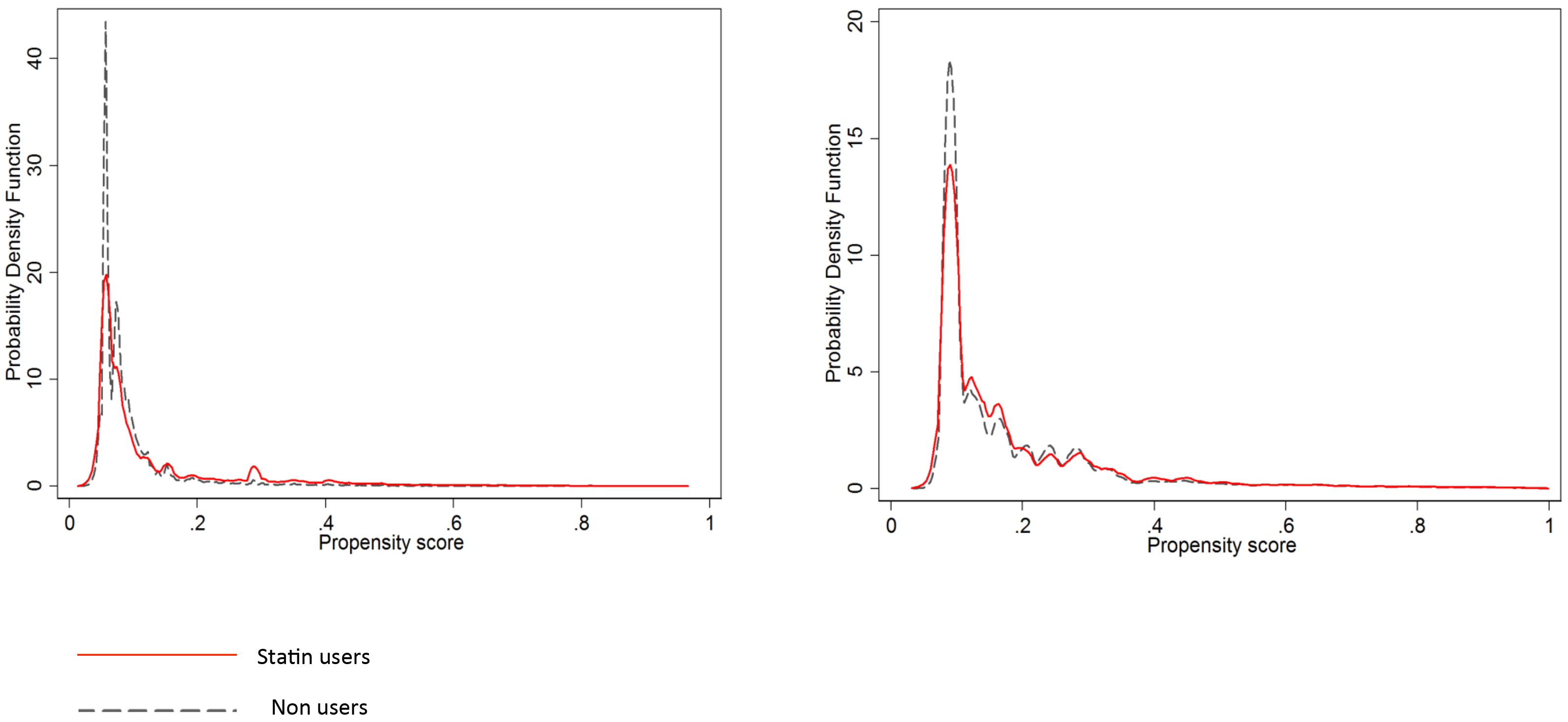
Distribution of the propensity scores (PS) among statin users and PS matched non-users in the initial PS matched population (n=268,087 (left)) and the final PS matched population (n=193,977 (right))

### Achieving clinical equipoise (matching step 3)

The results of matching steps 1 and 2 showed that the statin user and non-user groups remained unbalanced on the following baseline characteristics: myocardial infarct, congestive heart failure, peripheral vascular disease, cerebrovascular disease, diabetes, diabetes with end organ damage, prior use of low-dose ASA, and prior use of betablocking agents (See the column labelled “Initial PS matched study population” in Table 1). We therefore chose to match on these characteristics following the sex- and age matching (step 1) described above. This was done in the following manner: after the matching on sex and age, we initially matched on myocardial infarct. Subsequently, we matched on “remaining unbalanced morbidity” defined as having one of the following characteristics: congestive heart failure, peripheral vascular disease, cerebrovascular disease, diabetes or diabetes with end organ damage. Finally, we matched on “unbalanced pharmacological treatment”, defined as using either prior use of low-dose ASA or a betablocking agent. Subsequently, the propensity score matching (step 2) was repeated - however, without including the characteristics that were implemented in the clinical equipoise matching procedure described here (i.e. myocardial infarct, congestive heart failure, peripheral vascular disease, cerebrovascular disease, diabetes, diabetes with end organ damage, prior use of low-dose ASA and prior use of betablocking agents). This procedure resulted in balance on all covariates between the statin users and non-users (See the column labelled “Final PS matched study population” in Table 1 as well as Figure 2).

**Table 1.**
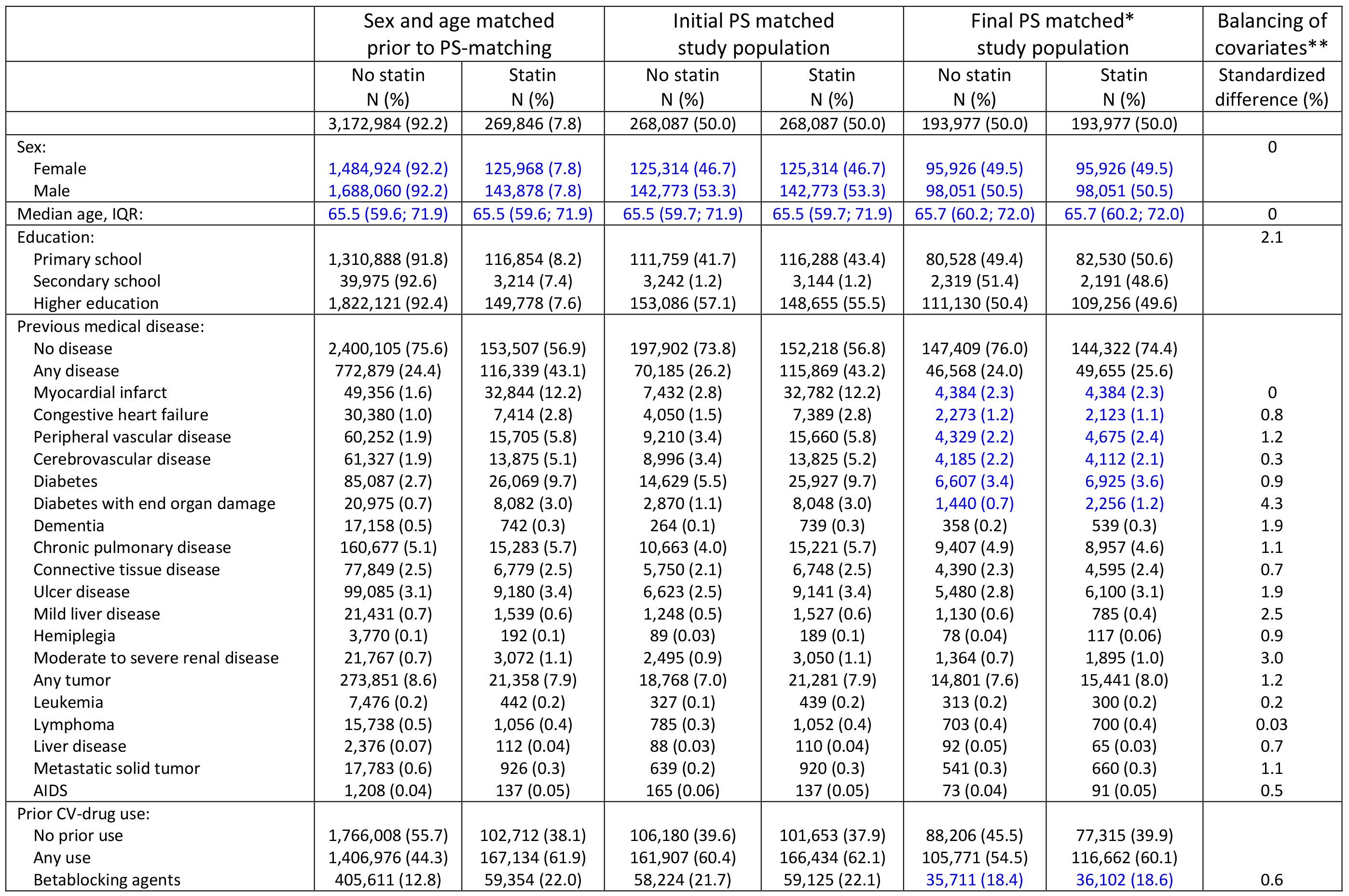
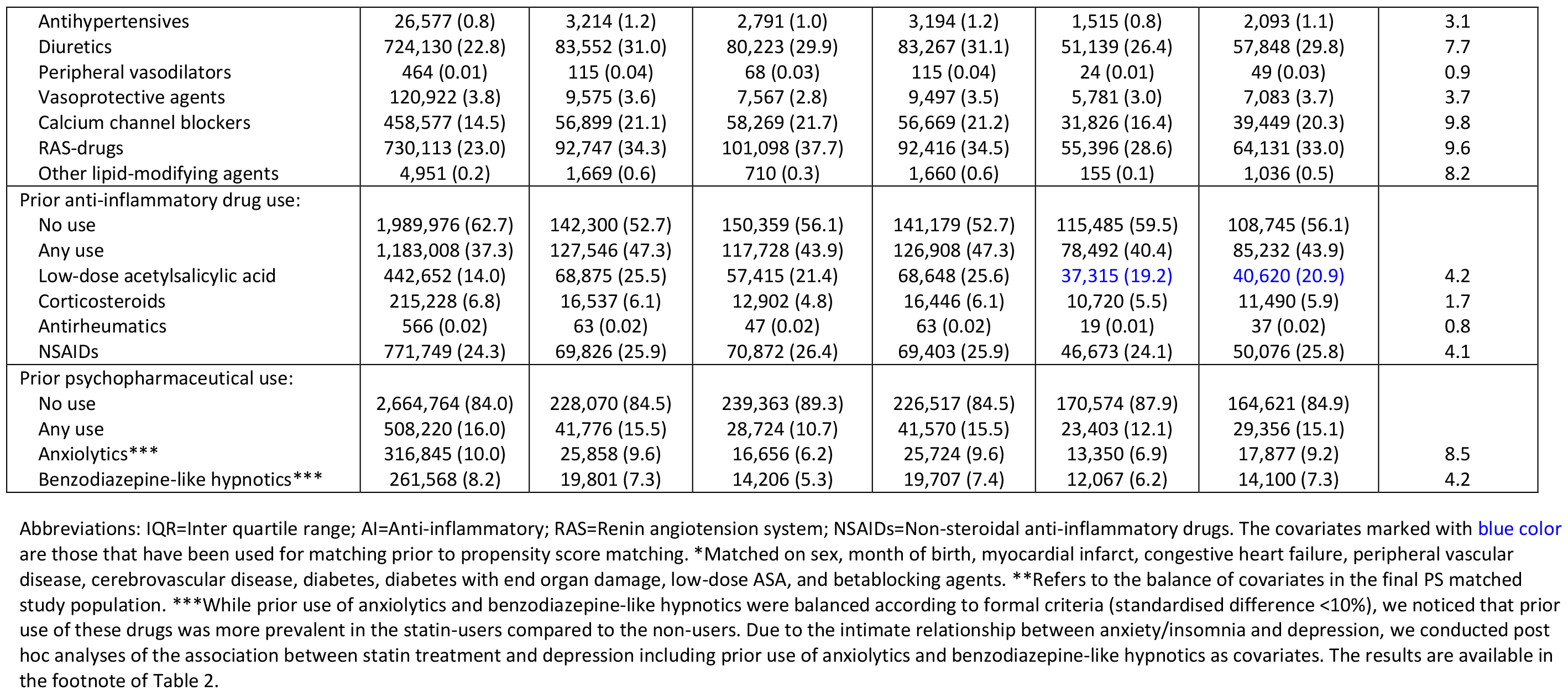
Baseline characteristics for the study population before and after propensity score (PS) matching

Due to this extra matching step, the final sample was reduced by 74,110 to 193,977 statin users and 193,977 non-users.

### Follow up

The statin users and non-users were followed from their index date until one of the following events: depression (see definition below under “Statistical analysis and outcomes”), cardiovascular disease, death, emigration or end of study period (December, 31, 2013). In accordance with the intent-to-treat approach, the statin users and non-users kept this treatment status from the index date through follow up. This approach gives a conservative estimate of the treatment effect by preserving the baseline comparability of the exposed and non-exposed groups, similar to that obtained in an intent-to-treat analysis of data from RCTs (35).

### Statistical analysis and outcomes

All analyses described below were carried out using STATA version 14.0. We used a stratified Cox regression model (allowing that each matched pair had their own underlying hazard in order to account for the matched nature of the propensity-score matched sample) with death as competing risk and calculated hazard rate ratios (HRR) with 95% confidence intervals (95%-CI) for the outcomes of interest. Depression was assessed in two ways: I) the first redemption of a prescription for an antidepressant (ATC-code: N06A) (23)and II) the first inpatient or outpatient diagnosis of depression (main ICD-10 diagnosis of F32 or F33) registered in the DCPRR (26). Since statins are known to decrease cardiovascular-and all-cause mortality in RCTs (1–5), we included the following “positive control” outcomes: I) the first diagnosis of cardiovascular disease (ICD-10: I00-99) based on data from the Danish National Patient Registry, II) cardiovascular mortality (ICD-10: I00-99) and III) all-cause mortality based on data from the Danish Cause of Death Register (36). The analyses of the five outcomes were carried out independently, i.e. the other outcomes (except from mortality) were not considered as competing risks.

In the first set of analyses, we compared the statin users to the matched non-users. Subsequently, we stratified the statin users on whether they received treatment as primary-or secondary prevention for cardiovascular disease. Secondary prevention was defined as having a diagnosis of myocardial infarct, congestive heart failure, peripheral vascular disease, cerebrovascular disease, diabetes or diabetes with end organ damage, i.e. the disease states that we matched upon in step 3.

### Post hoc analysis 1

Having observed the results from the initial analyses, showing a positive association between statin use and subsequent redemption of a prescription for an antidepressant (see Table 2), we speculated whether this association could be driven by either unmeasured confounding (despite the propensity score matching) or by a generally altered health care regimen initiated by the prescription of statins (control visits to general practitioner leading to more prescriptions in general etc.). To test this hypothesis, we assessed the association between statin use and redemption of prescriptions for any other drug than antidepressants and cardiovascular drugs (including statins) in the 193,977 statin users and 193,977 non-users using the exact same statistical approach as when antidepressant use was the outcome. The rationale being, that we would not expect the pharmacological effect of statins to cause an overall increase in morbidity that would elicit prescription of drugs in general (negative control). Thus, if the association between statin treatment and use of “any other drug” also turned out to be positive, it would suggest that the increased antidepressant use in those treated with statins would likely be due to either confounding, bias or “downstream” effects of the statin prescription-and not a causal (pharmacological) effect of the statins per se.

**Table 2.** Competing risk analyses of the association between statin use and the six outcomes of interest in the final propensity score matched study population (n=193,977 statin users and n=193,977 non-users) and stratified on individuals receiving statins for primary prevention (n=174,097 statin users and n=174,097 non-users) or secondary prevention (n=19,880 statin users and n=19,880 non-users) of cardiovascular disease, respectively.

|  | **N Total** | **N cases (%)** | **Rate per 10,000 person-years** | **HRR** | **95%-CI** |
| --- | --- | --- | --- | --- | --- |
| **Full sample** |  |  |  |  |  |
| **Antidepressant prescription<sup>1,\*</sup>** |  |  |  |  |  |
| Non users | 193,977 | 28,063 (14.5) | 215.4 | 1.0 | - |
| Statin users | 193,977 | 35,192 (18.1) | **279.3** | **1.33** | **1.31; 1.36** |
| **Prescription for any other drug<sup>2</sup>** |  |  |  |  |  |
| Non users | 193,977 | 186,248 (96.0) | **7,679.8** | 1.0 | - |
| Statin users | 193,977 | 190,870 (98.4) | **10,258.3** | **1.33** | **1.31; 1.35** |
| **Depression diagnosis<sup>3,\*</sup>** |  |  |  |  |  |
| Non users | 193,977 | 1,369 (0.7) | 9.6 | 1.0 | - |
| Statin users unadjusted | 193,977 | 1,629 (0.8) | 11.4 | **1.22** | **1.12; 1.32** |
| Statin users adjusted<sup>4</sup> | 193,977 | 1,629 (0.8) | 11.4 | 1.07 | 0.99; 1.15 |
| **Cardiovascular disease<sup>5</sup>** |  |  |  |  |  |
| Non users | 193,977 | 49,572 (25.6) | 279.3 | 1.0 | - |
| Statin users | 193,977 | 32,499 (16.7) | **247.0** | **0.89** | **0.88; 0.90** |
| **Cardiovascular mortality<sup>5</sup>** |  |  |  |  |  |
| Non users | 193,977 | 7,157 (3.7) | 58.8 | 1.0 | - |
| Statin users | 193,977 | 6,451 (3.3) | 54.2 | **0.92** | **0.87; 0.97** |
| **All-cause mortality** |  |  |  |  |  |
| Non users | 193,977 | 22,190 (11.4) | 182.4 | 1.0 | - |
| Statin users | 193,977 | 20,512 (10.6) | 172.3 | **0.90** | **0.88; 0.92** |
| **Primary prevention<sup>6</sup>** |  |  |  |  |  |
| **Antidepressant prescription<sup>1,\*</sup>** |  |  |  |  |  |
| Non users | 174,097 | 23,389 (13.5) | 204.0 | 1.0 | - |
| Statin users | 174,097 | 30,243 (17.4) | **273.4** | **1.36** | **1.34; 1.39** |
| **Prescription for any other drug<sup>2</sup>** |  |  |  |  |  |
| Non users | 174,097 | 166,612 (95.7) | 8,007.5 | 1.0 | - |
| Statin users | 174,097 | 171,092 (98.3) | **11,227.2** | **1.36** | **1.34; 1.39** |
| **Depression diagnosis<sup>3,\*</sup>** |  |  |  |  |  |
| Non users | 174,097 | 1,150 (0.7) | 9.2 | 1.0 | - |
| Statin users unadjusted | 174,097 | 1,409 (0.8) | **11.3** | **1.25** | **1.15; 1.36** |
| Statin users adjusted<sup>4</sup> | 174,097 | 1,409 (0.8) | **11.3** | **1.08** | **1.00; 1.18** |
| **Cardiovascular disease<sup>5</sup>** |  |  |  |  |  |
| Non users | 174,097 | 39,921 (22.9) | 242.4 | 1.0 | - |
| Statin users | 174,097 | 24,784 (14.2) | **212.1** | **0.90** | **0.89; 0.91** |
| **Cardiovascular mortality<sup>5</sup>** |  |  |  |  |  |
| Non users | 174,097 | 4,766 (2.7) | 43.8 | 1.0 | - |
| Statin users | 174,097 | 3,882 (2.2) | **36.7** | **0.92** | **0.89; 0.94** |
| **All-cause mortality** |  |  |  |  |  |
| Non users | 174,097 | 15,590 (9.0) | 143.3 | 1.0 | - |
| Statin users | 174,097 | 14,844 (8.5) | **140.2** | **0.94** | **0.92; 0.97** |
| **Secondary prevention<sup>6</sup>** |  |  |  |  |  |
| **Antidepressant prescription<sup>1,\*</sup>** |  |  |  |  |  |
| Non users | 19,880 | 4,674 (23.6) | 299.0 | 1.0 | - |
| Statin users | 19,880 | 4,949 (24.9) | **321.6** | **1.15** | **1.09; 1.21** |
| **Prescription for any other drug<sup>2</sup>** |  |  |  |  |  |
| Non users | 19,880 | 19,636 (98.8) | 5,698.7 | 1.0 | - |
| Statin users | 19,880 | 19,778 (99.5) | **6,679.6** | **1.25** | **1.20; 1.31** |
| **Depression diagnosis<sup>3,\*</sup>** |  |  |  |  |  |
| Non users | 19,880 | 219 (1.1) | 12.0 | 1.0 | - |
| Statin users unadjusted | 19,880 | 220 (1.1) | 12.1 | 1.03 | 0.82; 1.28 |
| Statin users adjusted<sup>4</sup> | 19,880 | 220 (1.1) | 12.1 | 0.96 | 0.77; 1.19 |
| **Cardiovascular disease<sup>5</sup>** |  |  |  |  |  |
| Non users | 19,880 | 9,651 (48.6) | 614.3 | 1.0 | - |
| Statin users | 19,880 | 7,715 (38.8) | **525.0** | **0.81** | **0.79; 0.83** |
| **Cardiovascular mortality<sup>5</sup>** |  |  |  |  |  |
| Non users | 19,880 | 2,569 (12.9) | 200.7 | 1.0 | - |
| Statin users | 19,880 | 2,391 (12.0) | **181.2** | **0.85** | **0.81; 0.90** |
| **All-cause mortality** |  |  |  |  |  |
| Non users | 19,880 | 6,600 (33.2) | 515.5 | 1.0 | - |
| Statin users | 19,880 | 5,668 (28.5) | **429.6** | **0.81** | **0.79; 0.84** |

Abbreviations: HRR = Hazard rate ratio; 95%-CI = 95% confidence interval. <sup>1</sup>Defined as the first redemption of a prescription for antidepressant medication (ATC-code: N06A). <sup>2</sup>Defined as the first redemption of a prescription for any drug except from cardiovascular drugs (including statins) or antidepressants. <sup>3</sup>Defined as the first depression diagnosis (ICD-10: F32-33) assigned at a psychiatric hospital. <sup>4</sup>The results on depression diagnosis are adjusted for antidepressant use during follow-up. <sup>5</sup>Defined by the ICD-10 codes for cardiovascular disease: I00-99. <sup>6</sup>Secondary prevention was defined as a prior diagnosis of myocardial infarct, congestive heart failure, peripheral vascular disease, cerebrovascular disease, diabetes, diabetes with end organ damage.
\*When adjusting the analyses of the association between statin treatment and antidepressant prescriptions for prior use of anxiolytics and benzodiazepine-like hypnotics, the HRR was reduced from 1.33 (1.31-1.36) to 1.29 (1.26-1.31) in the full sample, from 1.36 (1.34-1.34) to 1.30 (1.28-1.33) in the primary prevention group and from 1.15 (1.09-1.21) to 1.16 (1.10-1.22) in the secondary prevention group. When adjusting the analyses of the association between statin treatment and depression diagnosis for prior use of anxiolytics and benzodiazepine-like hypnotics (in addition to adjusting for antidepressant use following statin treatment), the HRR was reduced from 1.07 (0.99-1.15) to 1.05 (0.97-1.14) in the full sample, from 1.08 (1.00-1.18) to 1.06 (0.97-1.15) in the primary prevention group and from 0.96 (0.77-1.19) to 0.94 (0.74-1.18) in the secondary prevention group.

### Post hoc analysis 2

As the results from post hoc analysis 1 (there was a positive association between statin treatment and subsequent use of any other drug - with the exact same strength as the association between statin treatment and antidepressant use - see Table 2) indicated that the association between statin use and subsequent antidepressant use did not represent a causal effect, we investigated whether adjusting the analysis of the association between statin use and subsequent depression diagnoses assigned at psychiatric hospitals would be attenuated by adjusting for antidepressant use following statin use. If this turned out to be the case, the positive association between statin use and depression diagnoses would appear to be mediated by the use of antidepressants, which again seems to be due to either confounding, bias or downstream effects of the statin prescription-and thus not a causal (pharmacological) effect of the statins per se.

### Sensitivity analyses

The analyses outlined above were repeated after: I) stratifying on type of statin, and II) restricting the population to individuals born 1969-1983 in order to have complete data regarding prior psychiatric morbidity.

### Ethics

Use of the register data was approved by the Danish Data Protection Agency, the Danish National Board of Health and Statistics Denmark.

## RESULTS

The total follow up time for the 193,977 statin users and 193,977 non-users was 2,621,282 person-years (mean=6.8 years, SD=3.7). Among the statin users, 174,097 (90%) received statins as primary prevention and 19,880 (10%) as secondary prevention of cardiovascular disease. A total of 171,407 (88.4%) individuals used simvastatin, 12,658 (6.5%) used atorvastatin, 5,175 (2.7%) used pravastatin, 2,233 (1.2%) used fluvastatin, 1,067 (0.6%) used rosuvastatin, 803 (0.4%) used cerivastatin and 634 (0.3%) used lovastatin. Table 1 shows the baseline characteristics at the index date for the statin users and the matched non-users as well as the balancing of covariates (standardized difference) after successful propensity score matching.

Table 2 and Figure 3 show the main results of the study. Statin use was associated with increased risk of antidepressant use (hazard rate ratio (HRR)=1.33; 95% confidence interval (95%-CI)=1.31-1.36) as well as increased use of any other drug (HRR=1.33; 95%-CI=1.31-1.35 (post hoc analysis)). Statin use was associated with increased risk of receiving a depression diagnosis (HRR=1.22, 95%-CI=1.12-1.32), but not after adjusting for antidepressant use (HRR=1.07, 95%-CI=0.99-1.15 (post hoc analysis)). Statin use was associated with reduced cardiovascular disease (HRR=0.89, 95%-CI=0.88-0.90), reduced cardiovascular mortality (HRR=0.92, 95%-CI=0.87-0.97) and reduced all-cause mortality (HRR=0.90, 95%-CI=0.88-0.92). The findings of the analyses stratified on whether statin treatment was part of primary or secondary prevention of cardiovascular disease were similar to those reported above. The results of the sensitivity analyses (eTable 1 and eTable 2) are also largely consistent with those reported above.

**Figure 3.**
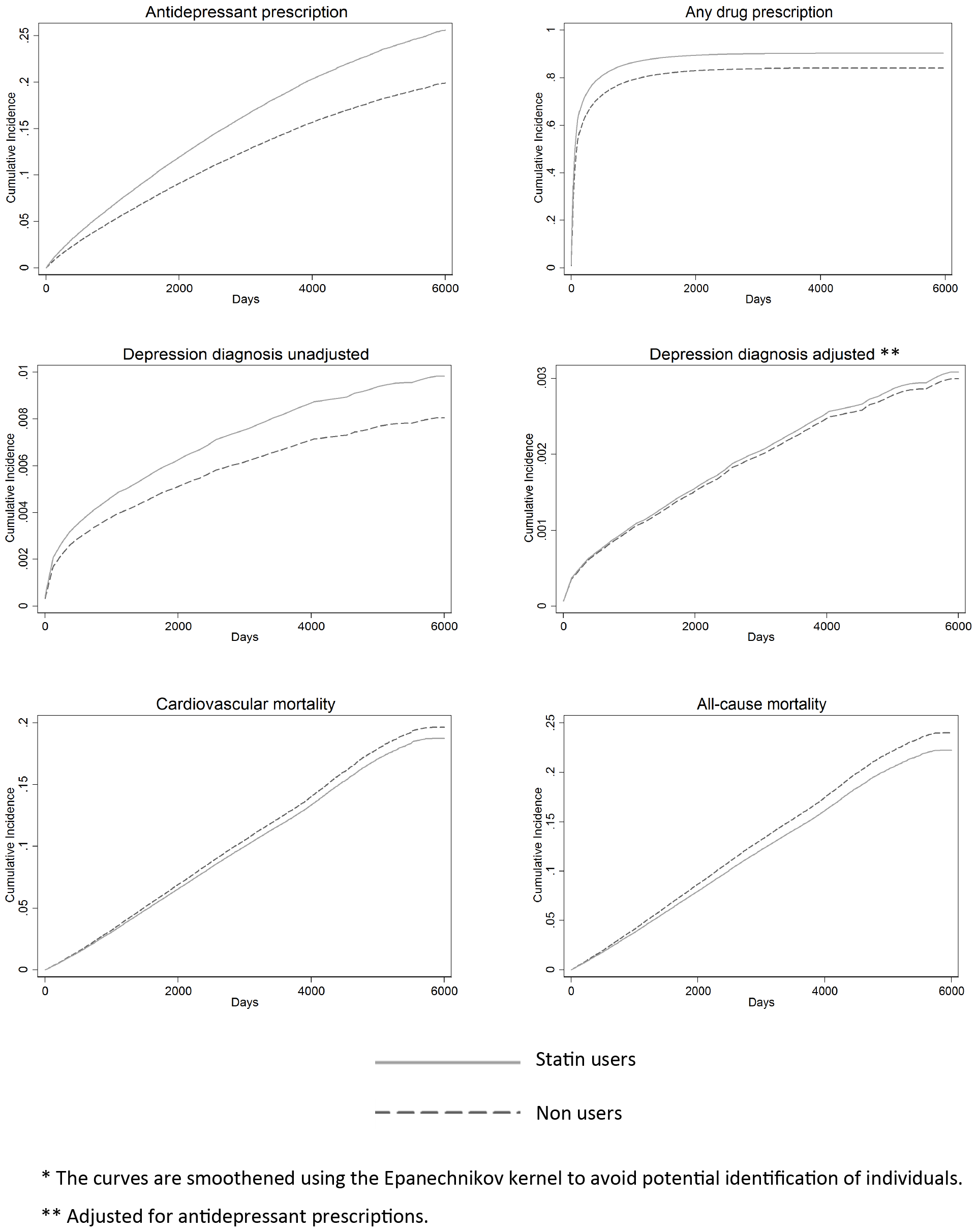
Competing risk analyses showing the cumulative incidence of redemption of a prescription for an antidepressant, redemption of a prescription for any other drug, receiving a depression diagnosis, cardiovascular mortality and all-cause mortality among 193,977 statin users and 193,977 propensity score matched non-users*

## DISCUSSION

In this study, 193,977 individuals treated with statins and 193,977 propensity score matched controls were followed for a total of 2,621,282 person-years. In the initial analyses, treatment with statins was associated with a significantly increased risk of antidepressant use and increased risk of receiving a depression diagnosis. However, the post hoc analyses showed that statin use was associated with a general/unspecific increase in prescription drug use. Hence, our initially detected link between statin treatment and depression is therefore highly likely to be driven by residual confounding or downstream effects of the statin prescription. Therefore, it appears that statin users and non-users are probably equally likely to develop depression, but that depression is simply more likely to be detected and treated among the statin users. The results regarding mortality support this interpretation of our findings. Had there been a substantial causal effect of statin use on the risk of depression, this is likely to have offset the beneficial effect of statins on mortality (via reduced cardiovascular morbidity/mortality) since depression is known to increase mortality (37). We observe the contrary, since the association between statin use and reduced allcause mortality was stronger than the association between statin use and reduced cardiovascular mortality (see Table 2). That statins reduced both cardiovascular mortality and all-cause mortality is in keeping with evidence from randomized controlled trials (1–5).

Our results indicating no causal association between statin use and the development of depression are consistent with those from the study by Mansi and colleagues who found no effect of statins on mental health (38), which, to our knowledge, represents the only prior propensity score matched observational study of the association between statin treatment and mental health. Our results are also in line with most, but not all (14) data stemming from RCTs on the effect of statin treatment on depressive symptoms in the context of prevention of cardiovascular disease (5,15).

Conducting large-scale placebo-controlled RCTs on the effects of statins on depression among people for whom statin treatment is indicated, i.e. in primary and secondary prevention of cardiovascular disease, is not ethically appropriate due to the well-proven effect of statins for these indications. Therefore, observational studies with proper handling of confounding, for instance via propensity score matching, represent the best alternative. We have conducted the largest propensity score matched study of the association between statin treatment and depression to date and the results do not support that statins have a causal effect on the risk of developing depression. On the contrary, the results indicate that statin users and non-users are probably equally likely to develop depression, but that depression is simply more likely to be detected and treated among the statin users.

## ACKNOWLEDGEMENTS

The authors are grateful to Katja G. Ingstrup, PhD (National Centre for Register-based Research, Aarhus University, Aarhus, Denmark) for her assistance with the propensity score matching.

## AUTHOR CONTRIBUTIONS

All authors contributed to the design of the study. Drs Köhler-Forsberg and Petersen conducted the analyses. The manuscript was drafted by Drs Köhler-Forsberg and Østergaard and was critically revised for important intellectual content by the remaining authors. The final version of the manuscript was approved by all authors prior to submission. Drs Köhler-Forsberg and Petersen had full access to all of the data in the study and take responsibility for the integrity of the data and the accuracy of the data analysis.

